# maxiM/Ze: An Image Recognition Approach for Visualizing and Processing Mass Spectrometry Based Metabolomics Data

**DOI:** 10.64898/2026.05.22.711157

**Authors:** Elizabeth R. Flammer, Timothy J. Garrett

**Affiliations:** Department of Chemistry, College of Liberal Arts and Sciences, University of Florida, Gainesville, FL; Department of Pathology, Immunology, and Laboratory Medicine, College of Medicine, University of Florida, Gainesville, FL

**Author notes:** Correspondence to: Dr. Timothy J. Garrett, Address: 1395 Center Dr. Room M641c, Gainesville, FL 32610.

## Abstract

Informatics is essential in metabolomics to analyze and interpret complex data for the advancement of biological insights. However, many current data-processing tools are time-consuming, require careful parameter selection, and depend heavily on user expertise, making reproducibility a challenge. To address these challenges, we developed maxiM/Ze, a Python-based application that utilizes image recognition algorithms to process liquid chromatography-high resolution mass spectrometry (LC-HRMS) metabolomics data prior to statistical analysis. The software implements an automated sequential pipeline that includes mass detection, extracted ion chromatogram (EIC) generation, peak alignment, and data visualization. By converting extracted ion chromatograms into PNG images, maxiM/Ze applies image processing techniques from OpenCV, including Canny edge detection, watershed segmentation, and Pearson correlation-based clustering, to align peaks across samples with minimal user input. Validation against Compound Discoverer 3.4 and mzmine 4.8.30 using eight replicate pooled plasma samples demonstrated competitive feature detection (12,067 features), annotation (219 unique compounds), and reproducibility (median CV of 35.8%) across platforms. The application is prepared for release on both Mac OS and Windows platforms, with the goal of improving reproducibility in metabolomics data analysis.

## INTRODUCTION

Informatics is essential in metabolomics to analyze and interpret complex data for the advancement of biological insights. Metabolomics via liquid chromatography-high resolution mass spectrometry (LC-HRMS) generates a large volume of data, necessitating computational approaches for analysis and interpretation. Prior to statistical analysis, raw data files from the instrument must undergo processing, which includes noise reduction, peak deconvolution, peak alignment, and metabolite annotation (1,2). Advanced informatics methods are required for statistical analysis, helping to identify patterns, correlations, and potential biomarkers using metabolomics data sets. This is crucial for understanding diseases and developing potential diagnostic and therapeutic strategies.

Although there are various open-source and commercial programs available for processing LC-HRMS files, the results are often heavily influenced by the chosen software and user expertise (3,4). Current software tools provide reliable outcomes for processing LC-HRMS data, however their limited interoperability across tools affects the comparison of results. In addition, these tools can be time-consuming, requiring hours to days depending on the study, and require careful selection of statistical parameters tailored to each unique dataset. This necessitates highly trained individuals in the fields of LC-MS and bioinformatics. As a result, reproducibility remains a challenge in metabolomics (5), and improper use of software can skew results leading to variability and false identifications (3).

To address these limitations, we propose maxiM/Ze, a software package that leverages image and pattern recognition algorithms to visualize and process LC-HRMS-based metabolomics data. By automating peak detection, alignment, and integration, maxiM/Ze minimizes human interaction to improve reproducibility and reduce processing time. The software is designed to scale effectively across large and varied datasets while remaining accessible through a user-friendly interface which lowers the barrier for researchers without extensive bioinformatics training. Additionally, maxiM/Ze was developed in accordance with the FAIR Principals for Research Software (6). Together, these features aim to advance the reliability and accessibility of metabolomics data analysis.

## METHODS

### Implementation

#### Pipeline

The pipeline imports all processing modules, initializes the file structure and mass groupings, and executes the full workflow for each mass group. Upon initialization, the pipeline loads file paths, configures mass groups from the configuration file, and assigns each mass a unique color for downstream visualization. The workflow is organized into checkpoints, each responsible for a specific processing stage: mass detection, extracted ion chromatogram (EIC) image generation, peak resolving, pixel mapping, peak slicing, coelution analysis, clustering, and visualization. Upon completion of all mass groups, the pipeline executes two final steps which includes exporting combined results (m/z, retention time (RT), peak height, and peak area) to an excel file, followed by optional library matching to annotate detected features if a library file was provided.

#### Mass Detection and Grouping

Users import their mass spectrometry data in mzML or mzXML format. At the time of import, users may specify predefined study groups, such as drug treated vs placebo. The software selects a representative set of six files, either randomly from the entire population or using a few from each predefined study group, for mass detection. This selection is saved to ensure reproducibility across repeated processing runs. Each file is scanned for centroid peaks exceeding a noise threshold of 5000 intensity units, and a peak must appear in at least seven consecutive scans to be considered a valid mass trace. Detected masses are grouped according to compatibility criteria. For the first grouping attempt, masses must differ by at least 5 ppm, must not have overlapping RT windows, must be separated by a minimum of one minute in RT, and must exhibit a similar intensity ratio of 90%. If masses are left ungrouped, the second grouping attempt relaxes the mass tolerance to 2 ppm and intensity ratio to 65% similarity. Detected features and their corresponding groups are exported to an excel file alongside summary statistics describing mass ranges, RTs, and intensities. A JSON file is also generated for downstream use within the pipeline.

#### Configuration

The configuration module stores the settings for the entire pipeline, including input and output directory paths, the analysis folder name, mass groupings, and tolerance parameters. Mass groups are loaded from the JSON file generated during mass detection and grouping.

#### Extracted Ion Chromatogram (EIC) Image Generation

This module reads mass spectrometry data files, generates EIC plots as PNG images (Figure 1), and exports a CSV of detected peaks for all masses within the current mass group. Each mass trace is smoothed using a Savitzky-Golay filter to improve peak detection. To be considered a peak the mass trace must rise above 4% of the maximum signal or three times the estimated noise level. Additionally, peak duration is constrained to approximately 0.03 to 0.75 minutes, and any peak exceeding 1.5 minutes is truncated. Peak shape quality is assessed by requiring at least a 25% rise and fall from the edges to the apex, with at least 70% upward slope preceding the apex and 70% downward trend following it. Finally, all output images are standardized to a consistent x-axis window of at least six minutes with 0.10 minutes of padding. Axis metadata and peak tables are saved to a CSV file.

**Figure 1.**
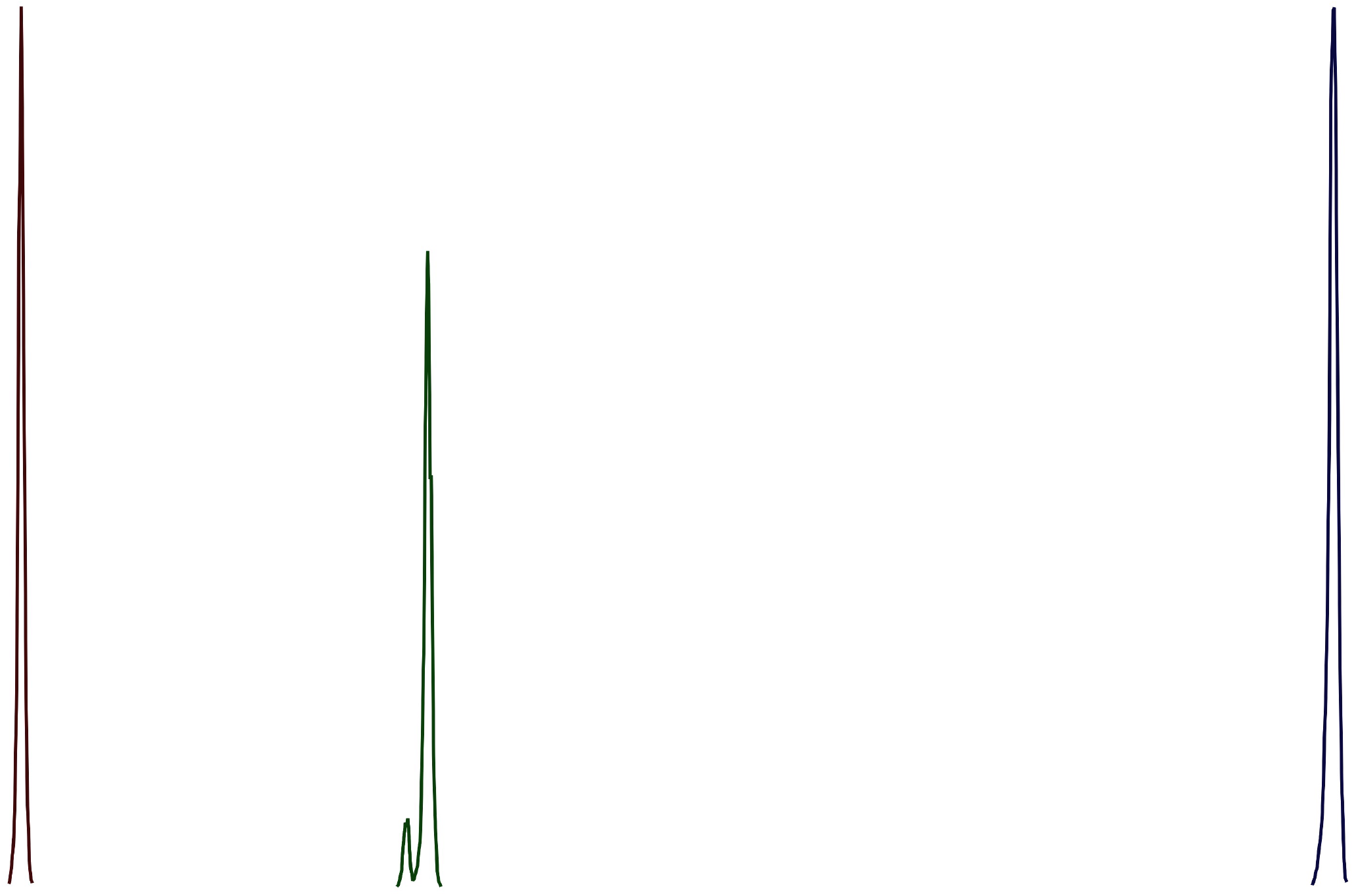
EIC PNG image generation output PNG at *m/z* values: 206.1005, 233.1383, and 393.2859.

#### Peak Resolving

The peak resolving module refines peak boundaries identified during EIC image generation using a combination of the peak CSV and EIC PNG. RTs are converted to x-axis pixel positions using the axis metadata saved for each PNG image. The module then evaluates coelution between peaks. Two peaks are considered resolved if their apexes are separated by at least 0.095 minutes and six pixels, and exhibit a valley drop of at least 40% between them. Peaks that do not meet these criteria are classified as coeluting. Additionally, peaks within 0.0355 minutes of a larger neighboring peak may alternatively be labeled as shoulder peaks rather than distinct coeluting features. To estimate peak boundaries, the module applies image processing tools from OpenCV, including Canny edge detection, distance transforms, and watershed segmentation, which estimates where one peak ends and another begins based on the image. A summary for each resolved peak is saved with updated RT start and end values, pixel start and end positions, cluster IDs, and peak type labels (resolved, coeluting, or shoulder). Additionally, the number of peaks per file is exported.

#### Peak to Pixel Mapping

Along with peak resolving, the peak to pixel mapping module utilizes the EIC PNG images to determine the exact pixel coordinates of each peak. For each image, the module identifies the colored traces corresponding to each mass and converts the visible trace regions into pixel start and end segments, which are saved to a CSV file. Each detected segment is recorded with the file name, *m/z* value, segment ID, pixel start, and pixel end, with segment IDs assigned sequentially from left to right across the image. The resulting CSV file provides exact horizontal pixel ranges for each colored mass trace.

#### Peak EIC Image Slicing

Each EIC PNG image is sliced into individual peak patch segments based on the pixel start and end positions determined during peak to pixel mapping (Figure 2A). Using the pixel mapping CSV file, the module crops the horizontal region corresponding to the pixel start and pixel end, while preserving the full image height. Every patch contains a single mass segment however, each patch may contain multiple peaks. Output patch segments are named using a combination of the sample name, m/z value, segment number, and group identifier.

**Figure 2.**
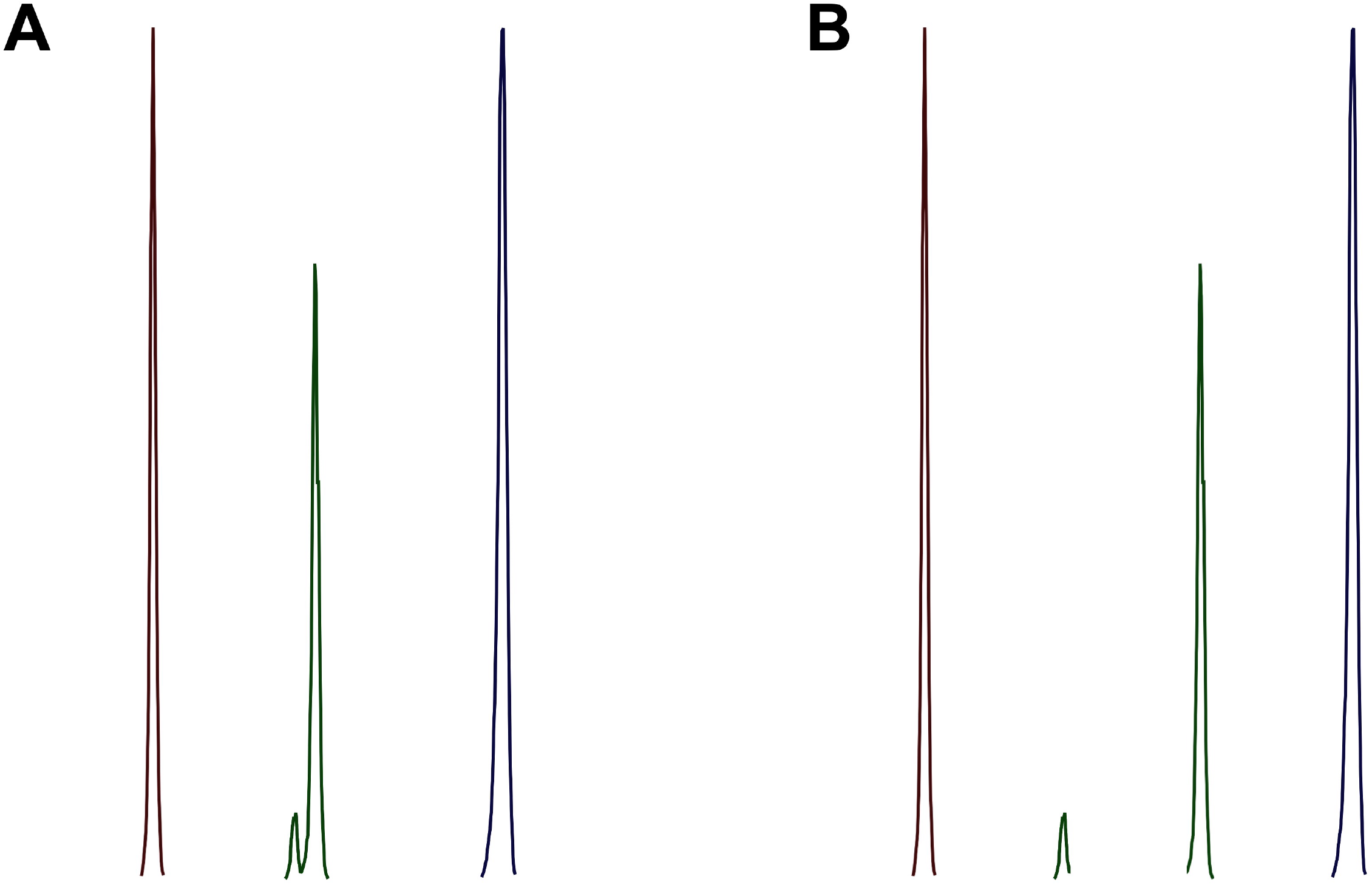
Slicing protocol for EIC peak patch generation. (A) Initial slicing of the EIC image based on peak to pixel mapping, resulting in three segment patches. (B) Additional slicing of a coeluting patch after coelution detection, resulting in four peak patches for clustering.

#### Peak Coelution

Peak patches containing coeluting peaks require special consideration, as coeluting peaks produce overlapping mass traces. This module determines which sliced peak images correspond to coeluting peaks and copied into a designated coelution folder, while non-coeluting slices are directed to a separate patch folder. Coeluting peaks are identified using the peak resolving and pixel mapping CSVs. Peaks that are labeled as coeluting in the peak resolving table are imported. Additionally, resolved peaks that share the same pixel mapping segment are reclassified as coeluting.

#### Slicing Coeluting Peaks

Coeluting peak patches identified in the previous step are further sliced into individual peak images (Figure 2B) by analyzing the intensity profile of each image and slicing at the lowest valleys between adjacent peaks. Each patch is first smoothed using a Savitzky-Golay filter. The expected number of peaks for each coeluting image is determined from the resolving CSV file. The image is then iteratively split at the lowest valley between adjacent peaks, and the resulting segments are saved. A warning is issued if the number of resulting patches does not match the expected peak count. Following slicing, the additional images are copied into the patch folder, and all patch images are renumbered to ensure every peak image has a standardized peak number to use for downstream processing.

#### Peak Clustering and Alignment

Following the slicing, a clustering and alignment step groups peaks across samples according to their m/z and peak shape by integrating PNG patch images with pixel coordinate data. Each patch image is converted to grayscale, resized to a standardized 50 by 50 pixel dimension, and smoothed with a 5 by 5 Gaussian kernel before being collapsed into a 1D intensity profile. Pairwise Pearson correlations between these profiles produce a square similarity matrix for all peaks sharing the same mass. A reference peak is then selected as the individual with the highest mean correlation to all others, serving as the most representative shape for that mass. Utilizing the reference peaks, peak patch images are clustered and aligned. The module produces a detailed peak alignment file, an alignment summary, a file that contains the unclustered patches that did not meet the criteria for any cluster group (if available), and a list of the patch image files within each clustered group.

#### Recluster Unclustered Peaks

Using the alignment summary and unclustered peaks list from the previous clustering step, the module attempts to place unclustered peaks into existing clusters based on mass and RT. For each missing sample value in the alignment table, it searches the unclustered peaks for a matching m/z and sample name, selects the candidate with the smallest RT difference within the defined RT tolerance window, and fills in the corresponding height and area values, marking the row as reclustered. The cluster PNG list is updated and the peak count recomputed, with each unclustered peak eligible for reclustering only once. Peaks that are successfully matched are removed from the unclustered list. The final output is an excel file containing three sheets (Summary, Unclustered, and Cluster PNGs), with newly filled cells highlighted in bold. This reclustering step recovers peaks missed during the initial clustering, due to subtle differences in peak shape, by leveraging m/z and RT based clustering.

#### Visualization of Peak Alignment

Following alignment and reclustering, two visualizations are generated. The first is a static PNG composite displaying peak regions across all samples, where each sample is assigned a unique semi-transparent color and each peak is a rectangle whose width corresponds to the RT range and the center is positioned at the peak apex RT. Dashed vertical lines indicate the aligned cluster centers for each mass and isomer position. The second is an interactive HTML composite where each clustered patch PNG image is placed along the RT axis at its aligned RT apex. Together, these visualizations allow users to assess whether the aligned peaks and cluster groupings are correct and consistent across all samples.

#### Final Export Excel Summary

Upon completion of all mass groups, the results from each group summary file are consolidated into a single excel file. The first sheet, *All_Group_Summary*, contains the group number, *m/z*, isomer position, aligned RT apex, peak count, reclustering status, and peak height and area for each sample file. The second sheet, *Groups_With_Unclustered_Peaks*, identifies any groups containing unassigned peaks, the number of unclustered peaks per group, and the file path to the corresponding group summary where each unclustered peak is listed.

#### Library Matching

To identify metabolites across features, detected features are matched against a reference compound library imported by the user. The top match is returned along with a set of secondary matches for each feature. Mass and RT tolerances for matching are configured by the user. The library file may be provided in CSV or excel format and must contain the columns “mz”, “rt”, and “name” but optional columns like “adduct” and “formula” are incorporated when available. The module produces a new excel file that appends library matches and corresponding columns to the original alignment summary, *All_Group_Summary*.

### Graphical User Interface

maxiM/Ze is slated for release for Windows (.exe) and macOS (.dmg) (Figure 3). Users import .mzXML or .mzML files with the option to define predefined study groups (e.g. Drug Treated, Placebo) from which samples are drawn during mass grouping or alternatively allow samples to be selected randomly. Target samples, such as pooled plasma controls, may also be included. Users then specify an output folder and run name, under which all results are saved. By default, mass groups are detected fresh from the input files each run; however, users may import a MassGroups_Cache.json from a previous run on the same sample set to bypass re-detection. An optional compound library (.csv or .xlsx) may be provided to match detected features against known compounds, with user-defined mass and retention time tolerances. Upon configuring all parameters, users initiate the pipeline, which performs peak detection, alignment, reclustering, visualization of peak alignment, and final Excel export of the alignment summary. Library matching results, when applicable, are appended to this summary as a separate output.

**Figure 3.**
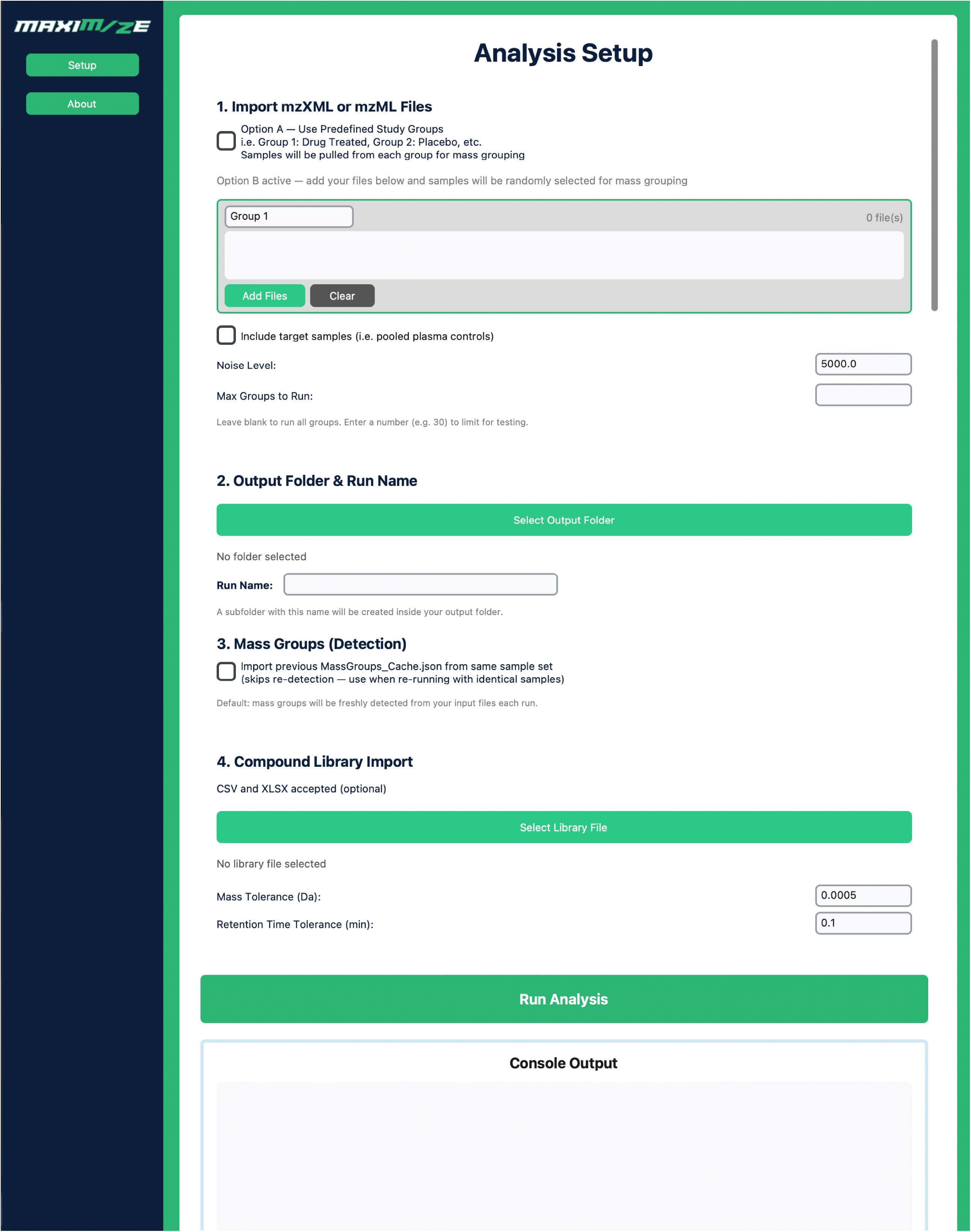
maxiM/Ze graphical user interface. The software dashboard displaying the parameter input fields.

## RESULTS

We analyzed eight replicate pooled plasma control samples obtained from the Red Cross using a Vanquish LC system coupled to an Orbitrap Exploris 120 operating in positive ionization mode, as previously described (7). Data processing was performed using maxiM/Ze, which detected 12,067 features across 2,064 mass groups, extracting both peak height and peak area for each. Features were matched against an in-house compound library containing 1,424 identifications (adducts counted as separate entries), yielding 279 matches, of which 219 were unique identifications and 60 were duplicates. Total analysis time was 28.6 hours.

maxiM/Ze was compared to Compound Discoverer (CD) 3.4 (Thermo) and mzmine 4.8.30 (mzio) for feature detection, annotation performance, and reproducibility across eight replicates. For feature detection, maxiM/Ze identified 12,067 features, falling between CD 3.4 (9,896 features) and mzmine 4.8.30 (16,173 features). Compound annotation yielded 144, 219, and 285 unique compounds for CD 3.4, maxiM/Ze, and mzmine 4.8.30, respectively (Figure 4), with 57 compounds confirmed across all three platforms and 181 shared exclusively between maxiM/Ze and mzmine 4.8.30. Notably, maxiM/Ze uniquely annotated 37 compounds not identified by either comparator. Duplicate compound identifications were observed across all platforms. Reproducibility, assessed by coefficient of variation (CV%) across replicates, was highest for CD 3.4 (median CV 23%), followed by maxiM/Ze (35.8%) and mzmine 4.8.30 (48.6%) (Figure 5). Analysis time varied considerably across platforms, with mzmine 4.8.30 completing in 56 seconds, CD 3.4 in 4.1 hours, and maxiM/Ze in 28.6 hours.

**Figure 4.**
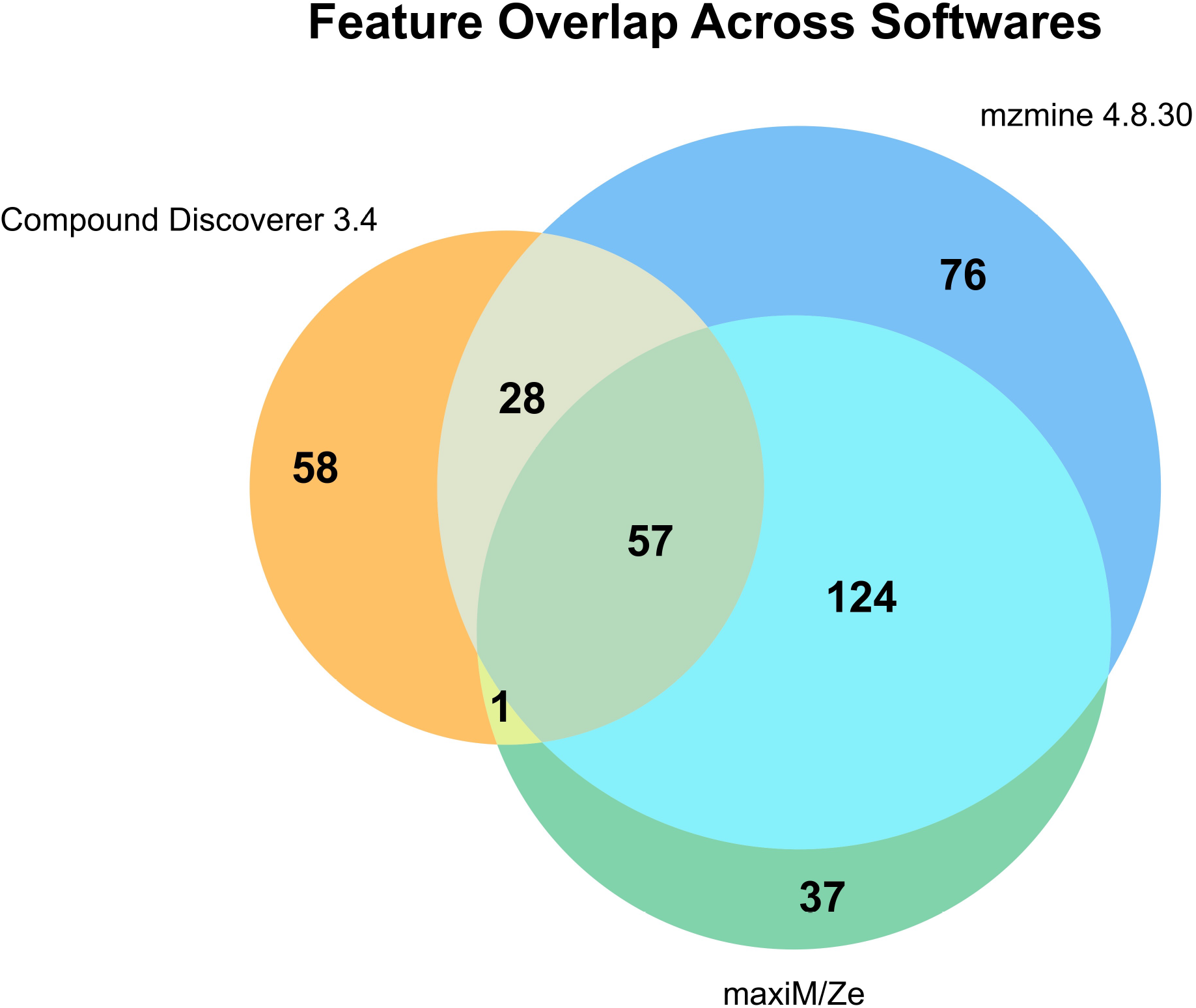
Venn diagram of unique and shared compound annotations across maxiM/Ze, CD 3.4, and mzmine 4.8.30. Annotations were performed against an in-house compound library, with 57 compounds confirmed across all three platforms and 181 shared exclusively between maxiM/Ze and mzmine 4.8.30.

**Figure 5.**
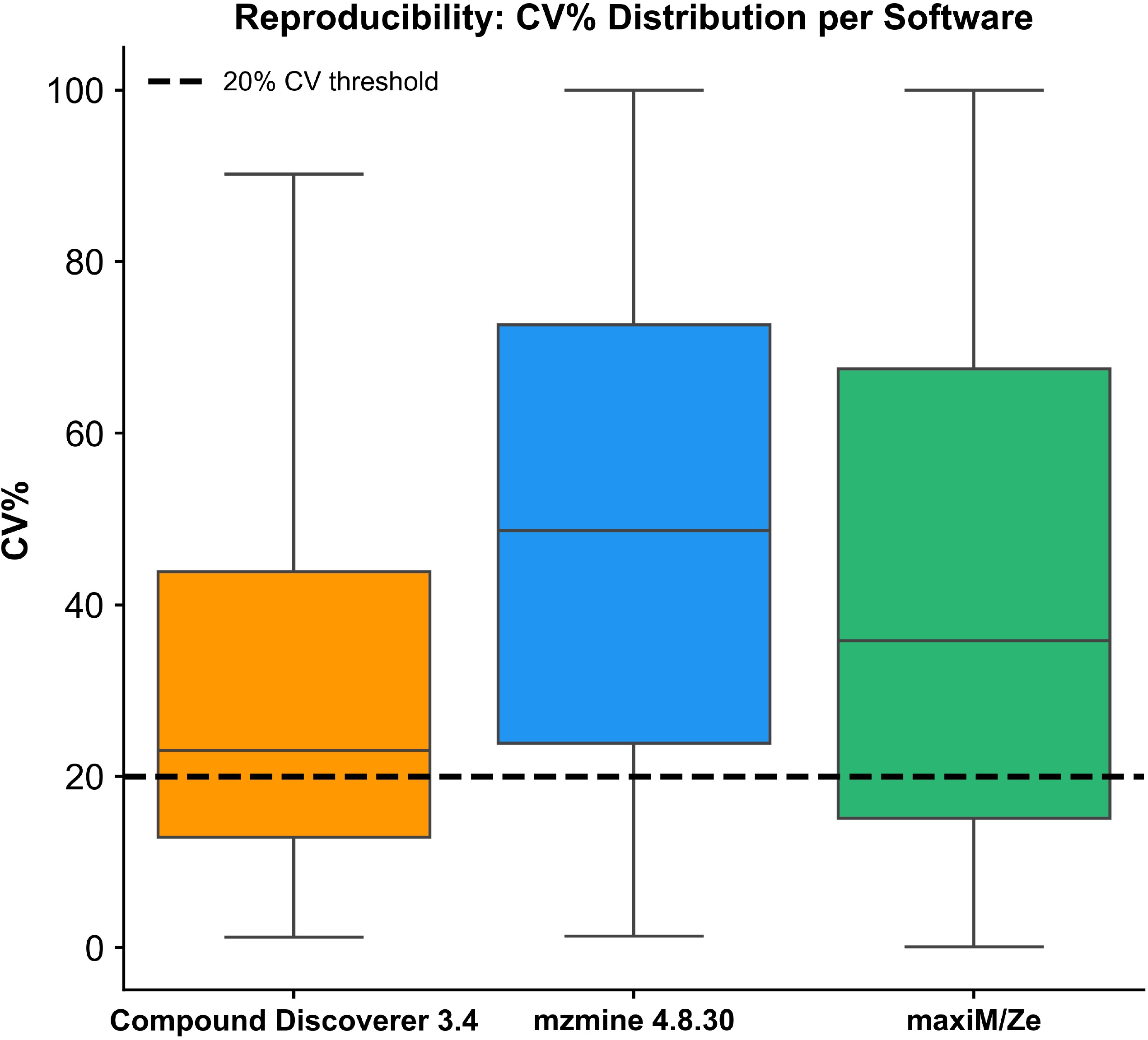
Box plot of CV% distributions across maxiM/Ze, CD 3.4, and mzmine 4.8.30 for eight replicate pooled plasma samples. The dashed line indicates the 20% reproducibility threshold. Median CV% values were 23%, 35.8%, and 48.6% for CD 3.4, maxiM/Ze, and mzmine 4.8.30, respectively.

## DISCUSSION

These results demonstrate that maxiM/Ze achieves a favorable balance between feature detection and measurement reproducibility. While CD 3.4 showed the lowest median CV%, its conservative feature detection and markedly lower annotation count (144 unique compounds) suggest a more stringent approach that sacrifices coverage. Conversely, mzmine 4.8.30’s expansive feature list was not accompanied by improved reproducibility, and its yield of high-quality features was comparable to maxiM/Ze despite detecting over 4,000 additional features. The strong annotation concordance between maxiM/Ze and mzmine 4.8.30 (181 shared compounds) provides external validation that maxiM/Ze reliably detects known metabolites.

Nevertheless, several limitations should be acknowledged. The overall median CV for maxiM/Ze (35.8%) remains above the conventional 20% reproducibility threshold, indicating that algorithmic refinements to peak integration or alignment may further improve performance. Analysis time also represents a current limitation, as maxiM/Ze’s 28.6-hour runtime far exceeds that of CD 3.4 (4.1 hours) and mzmine 4.8.30 (56 seconds), and remains an area of active development.

To further evaluate maxiM/Ze, we are currently assessing inter-user reproducibility across varying levels of expertise, including an undergraduate student, graduate student, and professor, by comparing results obtained independently using CD 3.4, mzmine 4.8.30, and maxiM/Ze.

## CONCLUSION

We have developed a Python-based application that processes LC-HRMS metabolomics data utilizing a novel image-based peak alignment approach and are preparing it for release on both Mac OS and Windows. Initial validation using eight replicate pooled plasma samples demonstrated that maxiM/Ze detected 12,067 features and yielded 219 unique compound annotations, performing competitively against both CD 3.4 and mzmine 4.8.30 in feature detection and annotation coverage. While CD 3.4 achieved superior reproducibility (median CV 23%), maxiM/Ze (35.8%) outperformed mzmine 4.8.30 (48.6%). Current limitations include a median CV above the conventional 20% threshold and a 28.6-hour analysis time, both of which are active areas of development. Our software is designed with a user-friendly interface that minimizes the learning curve for researchers, which is particularly important in multidisciplinary environments where users may not have extensive training in bioinformatics or mass spectrometry. Ongoing inter-user reproducibility testing across varying levels of expertise will further establish maxiM/Ze’s accessibility and reliability. By integrating advanced image and pattern recognition algorithms, we aim to improve peak detection and alignment, thereby increasing reproducibility and paving the way for more reliable research outcomes in metabolomics.

## ARTICLE INFORMATION SECTION

Author Contributions. E.R.F. developed and tested the software and wrote the manuscript. T.J.G. provided oversight of the research, reviewed all data and findings, and edited the manuscript. T.J.G. is the guarantor of this work and, as such, has full access to all the data in the study and takes responsibility for the integrity of the data and the accuracy of the data analysis.

